# Metagenomic analysis of soil samples collected from estuarine mangroves of Arabian Sea reveals rich microbiota and high numbers of sulphate reducing bacteria accompanied with methanogen bacteria

**DOI:** 10.1101/731810

**Authors:** Mandar S. Paingankar, Deepti D. Deobagkar

## Abstract

This study reports the analyses of the microbiome of the estuarine soil of mangroves of the Arabian Sea. Mangroves soil samples were collected from 12 locations of Arabian Sea coast of Maharashtra, India. 16S rRNA gene V3–V4 region amplicon sequencing was performed using the Miseq Illumina platform to identify the microbial communities present in the mangroves ecosystem. The metagenomics analysis provided an insight into the abundance, diversity and spatial variations in the mangrove microbial communities in relation to physico-chemical parameters and revealed that Proteobacteria, Flavobacteria and Planctomycetes are abundant in mangroves system. The differences in bacterial abundance, composition and diversity can partly be attributed to the physico-chemical characteristics of the samples, geographical location and anthropogenic activities in the locality. High numbers of sulphate reducing bacteria accompanied with methanogen bacteria were characteristic of Indian mangroves. The results obtained in the current study indicate rich species diversity and add valuable insights about the diversity of microbial communities of the mangroves in Maharashtra along the west coast of India and can provide better information for effective measures for conservation of mangroves. GIS based prediction suggest that the sulphur utilizing communities are under threat from anthropogenic activities and may decline in future if immediate measures are not implemented.

## Introduction

Mangroves system is known to play an important role maintaining the sea level and protecting the coast. These mangrove systems are characterized by structurally and functionally unique salt-tolerant, arboreal, flowering plant forests^1^. Taxonomically diverse group of plants and associated microbial communities are known to play fundamental roles in the nutrient cycle, productivity, functioning, and maintenance of the mangrove ecosystem^2^. The mangrove microbial community plays an integral role in nutrients cycling and thus nurturing the sustainable productivity^3^. Furthermore, fluctuations in temperature, aerobic conditions and salinity make these microbial communities more complex, diverse and unique^4^.Understanding the complex nature of mangrove microbial communities is one of the important aspects in microbes-plant interaction dynamics^3^. Similarly, microbes-plant interactions in mangroves have great implications in local and global ecological impact^5–8^. Furthermore these microbes are known to provide interesting biomolecules for human welfare^9^. Therefore exploration and identification of microbial communities in mangroves system have a potential application in human welfare as well as ecological conservation.

The microbial community studies have been carried out by various research groups to understand the role of microbial communities in the complex functioning of mangrove ecosystems^10–21^. In recent years, this ecosystem is rapidly getting destroyed and is threatened due to anthropological activities and modernisation^22–23^. In India, mangroves are present on eastern coastline, western coastline and in the Andaman and Nicobar Islands and are spread over an area of 4,921 sq. km. Presence of unique microbial communities in Sundarbans of India and Andaman Nicobar islands have been documented by various workers^12,13,16,18^. While on western coastline of India, mangroves are concentrated in Gujarat and Maharashtra. The mangroves of India are under immense pressure of conversion in to agriculture land and discharge of organic and inorganic matter from rapidly developing cities^9,23^. Mumbai, Thane, Raigad and Ratnagiri area in Maharashtra state are developing rapidly and anthropogenic activities in these are exerting the tremendous pressure on flora, fauna and microbial diversity^23,24^. Very few studies have been conducted to understand the microbial communities present in mangrove ecosystems of western coastline India^20,21,24^. Comparative studies on distinct mangrove microbial communities based on metagenomics will provide better insights into microbial community structure in the mangroves of India. Therefore the metagenomics explorative studies on microbial community structure in mangroves of India is essential in acquiring the baseline data on the diversity and functional genomics of microbial communities at higher resolution and in greater depth. The objective of the present study was to identify and to provide baseline data on the microbial community structure present in different mangrove areas along the west coast in Maharashtra State, India using the Illumina MiSeq sequencing platform.

## Material and Methods

### Ethics Statement

The collection sites of the current study are not part of any reserve forest or National park or privately-owned areas and therefore no specific permits were needed for the field studies. Endangered or protected species were not collected or included in the study.

### Sample Collection

Information on mangroves of India and shape files for GIS mapping were retrieved from earlier reports^22,23^. Sediment samples were collected superficially (0-5 cm depth) during the period of low tide at the GPS coordinates. Anjarle (17.840N, 73.101E and 17.843N, 73.103E), Kelashi(17.928N,73.074E and 17.931N, 73.077E), Bankot(17.958N, 73.031E and 17.961N,73.033E), Uran(18.896,72N.939E and 18.898N,72.944E), Agarkot(18.546N,72.934E and 18.549N,72.943E), Vashim(18.812N,73.033E and 18.814N,73.026E). These mangrove forests were located in estuarian regions of the Jog, Bharja, Savitri, Kundalika and Patalgangarivers of Maharashtra, India (Supplementary Information Table S1).

### Environmental parameters

The average temperature, average rainfall, pH, SO4, NO3, PO4, Mg, Cl2 and salinity of samples was measured and is depicted in Supplementary information Table S2.

### DNA extraction from soil samples

Soil collected from respective samples was processed for DNA isolation using Powersoil DNA isolation kit (MoBio laboratories Inc. Carlsbad, CA) as per manufacturer’s instructions. DNA concentration was measured using the Quantus fluorimeter (Promega, USA).

### Amplification primers

Regions corresponding to V3 and V4 regions of 16s rRNA were amplified from the DNA samples and were subjected to pyrosequencing using the Miseq Illumina platform.

### Sequence analysis

Sequence analysis was carried out by employing the methods described earlier^24^. The taxonomic assignment of unassembled metagenomic sequences was performed using BLASTX against the SEED and Pfam databases on the MG-RAST server v2.0 (http://metagenomics.nmpdr.org) using a cut-off E-value of 1e-10. BLASTX was also used to conduct a similarity search against the NCBI-NR database, and MetaGenome Analyzer software (MEGAN v6.0)^25^(https://github.com/danielhuson/megan-ce) with the LCA algorithm (maximum number of matches per read: 5, min support: 5, min score: 35, top percent:10) was used to visualize results.

### Statistical analysis

Several indices of clonal diversity were estimated using the MEGAN and PAST3 program available from the University of Oslo website link (http://folk.uio.no/ohammer/past). A similarity matrix was generated using a probabilistic distance metric. The statistical significance of the relationship between the differences in chemical composition and (i) species diversity, (ii) species alpha diversity indices and (iii) species beta diversity indices was tested by Mantel tests. P values were calculated using 9999 permutations on rows and columns of dissimilarity matrices.

### GIS mapping and prediction using DIVA GIS and MAXent

The mangroves sulphur utilizing bacteria data was retrieved from current study and earlier study^24^. The baseline (1950–2000) temperature and precipitation layers were obtained from WorldClim-Global Climate data repository (www.worldclim.org). Temperature and precipitation layers of the future (year 2020, year 2030) were obtained from repository (www.ccafs-climate.org). The data were resampled at the native WorldClim 30 arcsec (approximately 1×1km) resolution. Statistical modelling was conducted with DIVA GIS and MAXENT (https://biodiversityinformatics.amnh.org/open_source/maxent/) software with the present data and from the environmental variables.

## Results

### Description of the Community

Sequence clustering resulted in the identification of 1357 (364.16±84.80 per sample) different bacterial species (Supplementary Information Table S3). In general, Proteobacteria and Bacteroidetes were more abundant and other taxon such as Cyanobacteria, Euryrchaeota, Chloroflexi and Verrucomicrobia were present at moderate abundance. The Acidobacteria were more abundant in Kelashi sample whereas Fusobacteria showed moderate abundance in the Uran sample. The numbers of sequences affiliated with each taxon are depicted in Figure 2A, with a major abundance of Gammaproteobacteria (8.86-20.26 %), Flavobacteriia (7.24-25.76%), Alphaproteobacteria (9.34-24.65%), Deltaproteobacteria (2.88-14.57%) and Actinobacteria (0.17-6.1%) followed by other minor class represented by Planctomycetia (1.2-3.57%). Sulphate utilizer species such as *Desulfobulbus mediterraneus, Desulfomicrobium sp., Desulfonatronum thiosulfatophilum, Desulfosarcina sp.* were present in most of the samples. These sulphate reducers were accompanied by methanogens at all sites. *Aciditerrimonas ferrireducens, Algisphaera agarilytica, Ardenticatena maritima, Bacteriovorax marinus, Bdellovibriobacteriovorus, Bellilinea caldifistulae, Blastocatella fastidiosa, Blastopirellula marina, Calothrixdesertica, Candidatus Solibacterusitatus, Litorilineaaerophila, Ornatilinea apprima, Owenweeksia hongkongensis, Phycisphaera mikurensis Thiohalobacterthiocyanaticus Trueperaradiovictrix* were present in moderate numbers in all collected samples. *Deferrisomacamini* exclusively present in ASUN samples whereas *Nafulsella turpanensis Roseiflexus sp*. RS-1 exclusively present in KLM samples. Species such as *Lewinella nigricans Methanomassiliicoccus luminyensis, Methanothermococcuso kinawensis, Robiginitaleabiformata* showed moderate presence in all sampling sites except KLM samples. *Ignavibacterium album* and *Ilumatobacter fluminis* were present in moderate numbers only in ASUN, ASWD, BNM samples whereas *Gramella aestuarii* present significant number only in ADM BNM samples.

**Table 1:** Details of soil samples collected from mangroves

| Place | District | Sample code | Latitude | Longitude | River |
| --- | --- | --- | --- | --- | --- |
| Anjarle | Ratnagiri | ADM1 | 17.840 | 73.101 | Jog River |
| Anjarle | Ratnagiri | ADM2 | 17.843 | 73.103 | Jog River |
| Kelashi | Ratnagiri | KLM1 | 17.928 | 73.074 | Bharja River |
| Kelashi | Ratnagiri | KLM2 | 17.931 | 73.077 | Bharja River |
| Bankot | Ratnagiri | BNMN1 | 17.958 | 73.031 | Savitri River |
| Bankot | Ratnagiri | BNMN2 | 17.961 | 73.033 | Savitri River |
| Uran | Raigad | ASUN1 | 18.896 | 72.939 |  |
| Uran | Raigad | ASUN2 | 18.898 | 72.944 |  |
| Agarkot | Raigad | ASWD1 | 18.546 | 72.934 | Kundalika River |
| Agarkot | Raigad | ASWD2 | 18.549 | 72.943 | Kundalika River |
| Vashim | Raigad | VASH1 | 18.812 | 73.033 | Patalganga River |
| Vashim | Raigad | VASH2 | 18.814 | 73.026 | Patalganga River |

**Figure 1:**
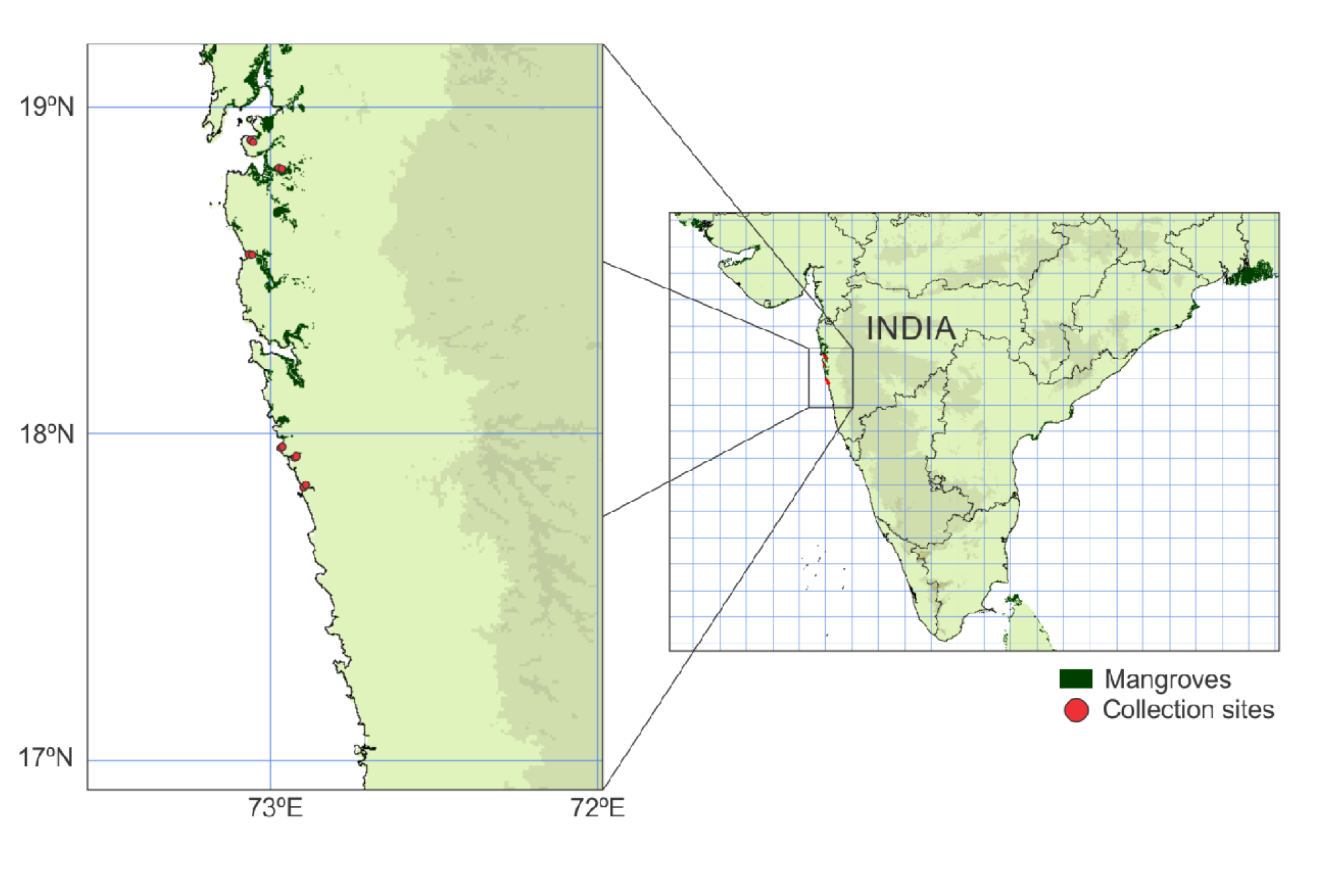
Sampling sites: Mangrove soil samples were collected from different locations from eusturine mangrove along the coast of Arabian Sea.

**Fig. 2.**
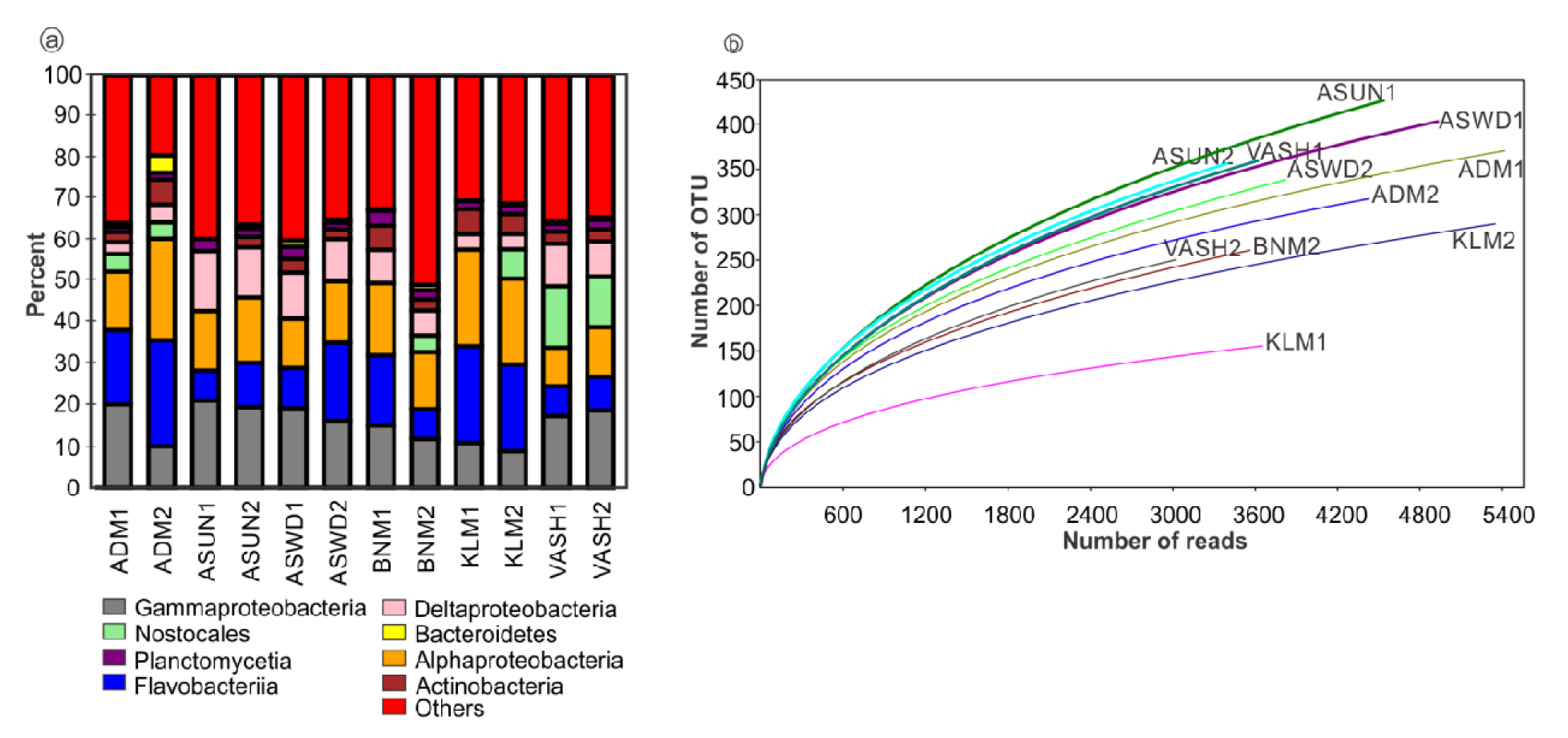
Distribution of predominant bacterial class in samples based on 16S rRNA gene sequencing. **A)** Class distribution Observations are displayed as stacked bar charts for individual mangrove sample (x-axis) against the percent class abundance (y-axis). B) Rarefaction curves for mangrove samples

### Alpha and Beta diversity of bacterial communities in different mangrove soil samples

Alpha and beta diversity analysis revealed rich taxonomic diversity and dominance of few species in mangroves samples (Supplementary Information Table S4; S5). Interestingly, soil samples from KLM1 (D = 0.06438), KLM2 (D = 0.03303),VASH1 (D = 0.03259) and VASH2 (D = 0.03802) showed higher dominance of fewer bacterial groups. Chao-1 analysis identified (274-729) species in each sample. At species level, high beta diversity was observed in all mangrove samples (Supplementary Information Table S5) wherein 36 species were common in all 12 samples. ASUN1 sample showed maximum number (61) of unique species.

### Habitat Type Differences in Communities

In order to examine the community structure and its specific features, PCoA analysis was carried out. Most samples clustered closely together, indicating that the microbial communities residing in these mangroves are similar and represent characteristic and typical community structures. Interestingly, communities in the mangroves facing industrialization threats, KLM1, KLM2 and ADM1, ADM 2 were different from all other (Fig. 3A). Similarly, neighbour joining tree revealed the distinctness of KLM1, KLM2 and ADM1, ADM 2 samples (Fig. 3B). Canonical correlation analysis (CCorA) was employed to find out the correlations between alpha diversity indices and environmental factors. A strong positive correlation was observed between the SO4 concentration and relative dominance of a few species. A positive correlation was observed in PO_4_ and Cl_2_concentration and Berger-Parker index in samples. A negative correlation was observed in NO_3_concentration and dominance in samples(Fig. 4A). Mantel test was used to determine the factors (diversity, chemical parameters, and Whitaker beta diversity) which best predicted the community diversity across the different mangrove areas samples (Fig. 4B). Differences in chemical parameters correlated significantly with diversity across different collection sites (r = 0.278; p value = 0.02). Similarly, chemical parameters of soil had significant effect on diversity and beta diversity (r = 0.717; p value = 0.0001). These results indicated that soil chemical composition and presence of pollutants is playing an important role and influences the bacterial diversity.

**Figure 3:**
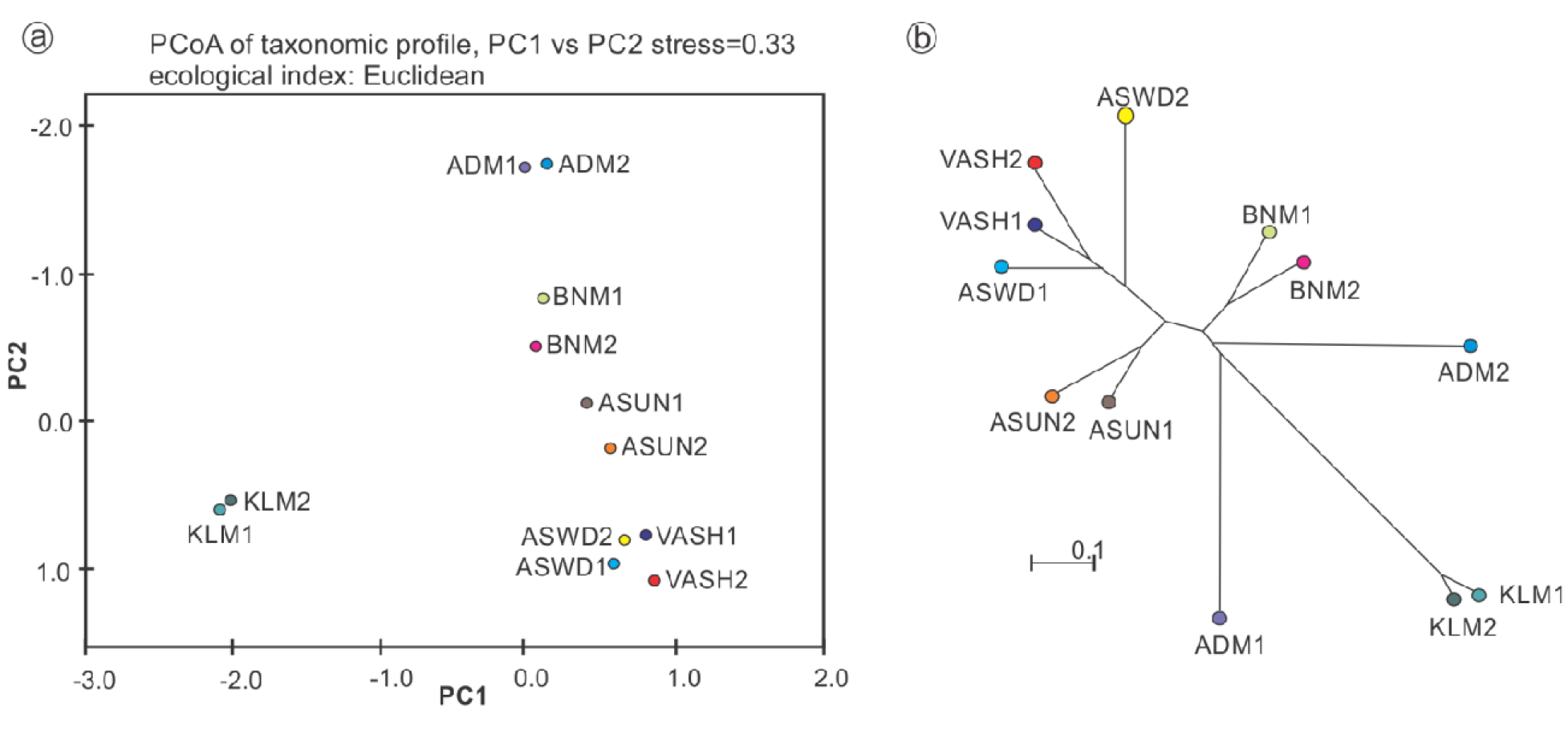
A) Principal coordinate analysis (PCoA) of the bacterial communities derived from the weighted UniFrac distance matrix. B) Neighbor-joining phylogenetic tree

**Figure 4:**
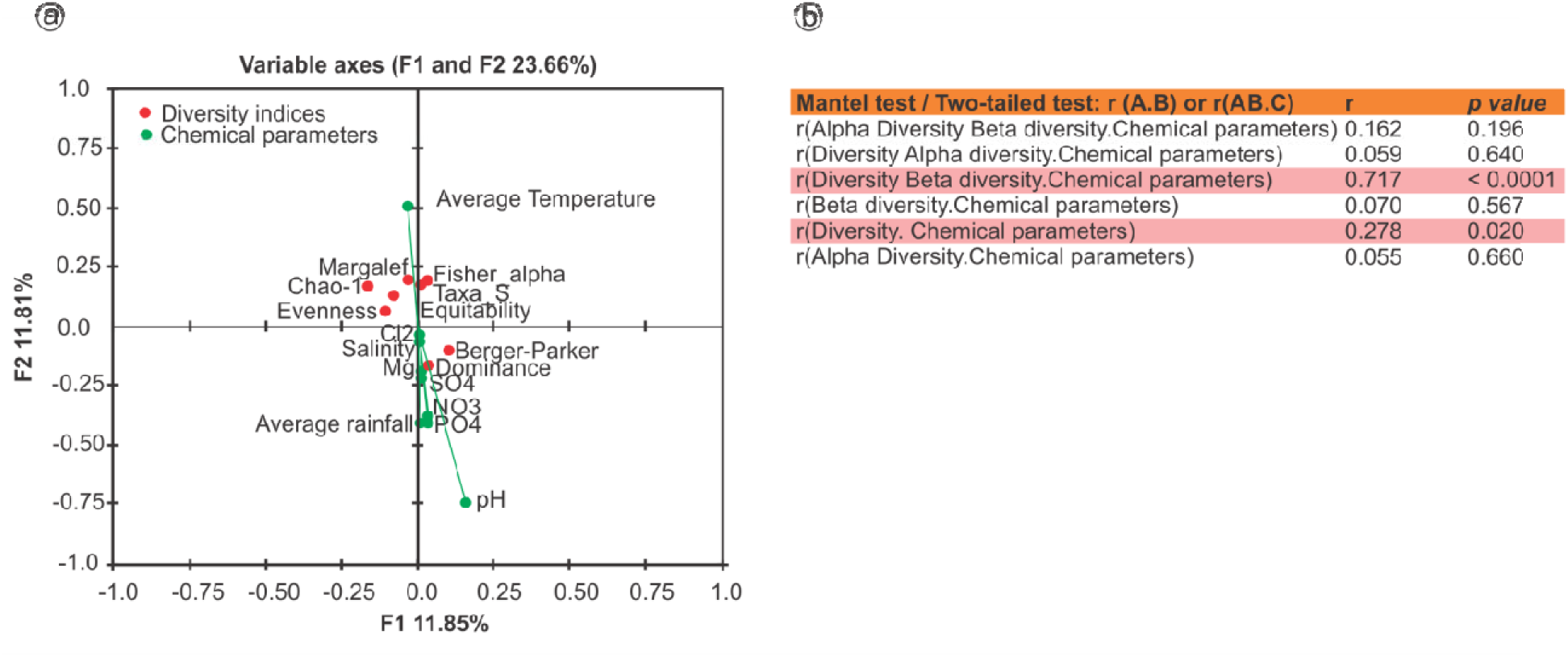
A Canonical correlation analysis (CCorA) B) Mantel Test

### GIS based prediction of distribution of sulphur utilizing bacteria in Maharashtra

Using the current climate layers (68 layers) the distribution of sulphur utilizing bacteria in Maharashtra was predicted (Fig. 5). The prediction suggests that areas such as Dighi, Chiplun, Uran, Jawaharlal Nehru port are suitable for growth of sulphur utilizing bacteria. Based on future climatic conditions, sulphur bacteria prediction map for the year 2020 and 2030 was predicted. The prediction suggests that area suitable for growth of sulphur utilizing bacteria will be decline in 2020 and 2030 (Fig. 5).

**Figure 5:**
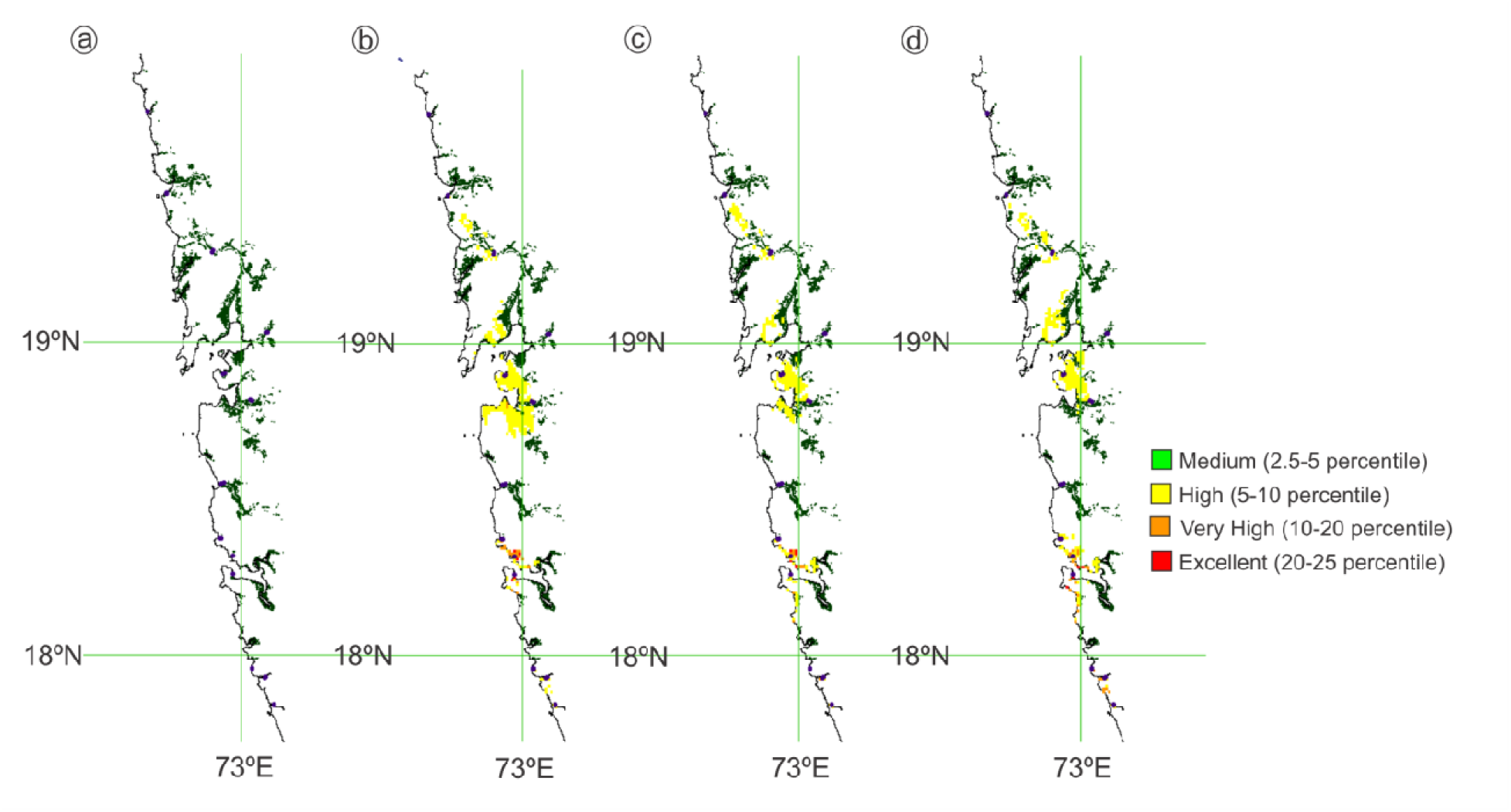
GIS based prediction of sulphur utilizing bacteria in Maharashtra

**Figure 6:**
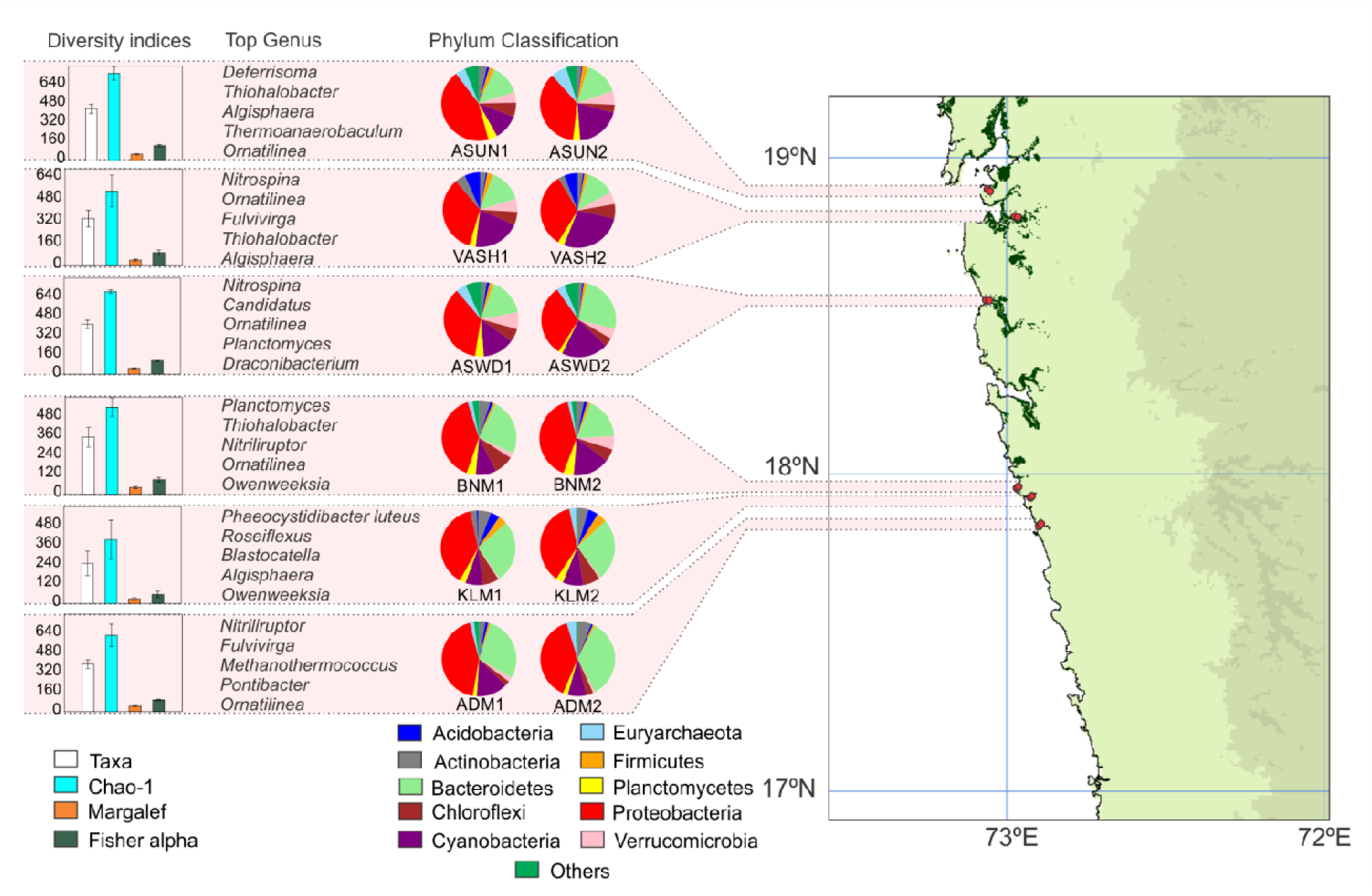
Microbial diversity at various mangrove sites

## Discussion

The mangroves of India are under tremendous pressure of anthropogenic activities and are rapidly declining^23^. Maharashtra state of the west coast of India is one of the rapidly industrializing and urbanizing region and discharging a huge amount of industrial and organic waste to Arabian Sea^26,27^. Jha Fernandes. The draining of industrial and domestic waste the mangroves of Maharshtra is significantly affecting the mangroves and associated flora, fauna and microbes. Therefore there is urgent need to gather the baseline data of microbes associated with mangroves as they play important role in mangroves ecosystem dynamics. In the current study metagenomic profiles of mangrove sediments of west coastline of India was carried out using the miSeq Illumina platform. All the samples showed the high abundance of bacterial reads and a small percentage of sequences from Archaea and unidentified organisms.

A taxonomic analysis of samples collected from Kerala, Tamil Nadu, Sundarbans, Andaman Nicobar islands, Maharashtra and current study clearly indicates the higher abundance of Proteobacteria in the mangroves sediments^12,16,17,18,20,21,24^ (Fig. 2A). The metagenomic analysis identified more than 80% species inhabited in these mangrove sediments (Fig.2B). Among the Proteobacteria detected, Gammaproteobacteria were most abundant followed by Alphaproteobacteria. The nifH gene, the key players in nitrogen fixation, has been detected predominantly Alphaproteobacteria, Betaproteobacteria, and Gammaproteobacteria^30–32^(Fig. 2A).Along with Proteobacteria, *Bacteroidetes*, *Acidobacteria*, *Firmicutes*, *Actinobacteria*, *Nitrospirae*, *Cyanobacteria*, and *Planctomycetes* were also showed their moderate presence in the current study and Indian mangroves^12,16,17,18,20,21,24^. A similar patterns were obtained from the samples belonging to mangroves areas of different parts of the world^11,14,15,19,20,21,28,29^. As compared to Indian and Brazilian mangrove samples, Red sea mangrove samples showed relatively higher number of Achaea^12,16–21,24^. Bacteroidetes known for hydrocarbon degradation and substantial abundance of Bacteroidetes has been previously noted from mangroves samples^10,16,20,21,24,33^.The mixed marine and terrestrial conditions in mangroves are suitable for nitrogen fixer proteobacteria and Bacteroidetes. Current study and samples collected from Kerala showed the moderate numbers of Actinobacteria^20,21^. Although Chloroflexi and Verrucomicrobia detected routinely in metagenomics analysis, little is known about the possible role of these bacterial phyla. Samples from Indian mangroves showed the presence of high numbers of sulphate reducing bacteriaaccompanied with methanogen bacteria. Sulphate reducing bacteria, such as *Desulfobulbus, Desulfomicrobium, Desulfonatronum, Desulfosarcina*were found accompanied with methanogen genera, including *Methanothermobacter, Methanococcus, Methanocaldococcus, Methanosarcina, Methanococcoides, Methanospirillum*^12,16,17,18,20,21,24^. The results obtained in current study and earlier studies clearly suggest that sulphate reducers and methanogens might be using the different metabolic pathways in non-competitive manner to shape up the microbial metabolism in mangroves ecosystem^34–38^. Sequence reads assign to aromatic hydrocarbons degraders and an iron and sulphur-reducing mesophilic anaerobe were found significantly lower in Indian mangroves samples.

The alpha and beta diversity indices clearly indicate the presence of high species richness and high diversity in these samples. PCoA and neighbour joining analysis clearly suggest the distinctness of KLM and ADM samples from rest of the samples (Fig. 3A,3B). Relatively less organic and inorganic pollution might be responsible for distinctness of KLM and ADM samples. The Canonical correlation analysis and Mantel test clearly suggest that chemical properties of soil sediments and micro environment in this ecological niche influenced bacterial diversity in these mangrove areas (Fig. 4A;4B). It has been well documented that geographical location, plant species, and/or physico-chemical parameters play an important role in shaping the microbial structure. The observed variation in microbial diversity and composition in studied samples might be attributed to the environmental factors and anthropogenic activities in respective mangrove forests.

GIS based prediction tools are important in examining the potential problems arising from natural and artificial forces on environment as well as microhabitats. Furthermore, GIS is one of the important tools which can be used in ecology to preserve and monitor the makeup of biological life within a given area. However, its use in predicting microbial ecology is limited^39,40^. In the current study GIS prediction was used to predict the sulphur utilizing bacteria in Maharashtra region. Similarly future climate data was used to predict the trends of sulphur utilizing bacteria in response to climate change. Using current climate condition layers (total 68 layers) the distribution of sulphur utilizing bacteria in Maharashtra was predicted. The prediction suggests that climatic conditions in Raigad district especially near Dighi, Uran and Jawaharlal Nehru port are suitable for growth of sulphur utilizing bacteria. These bacteria are useful in recycling of various compounds such as sulphates, alkanes and many more. The prediction using future data suggest the decrease in suitable area for growth of these sulphur utilizing bacteria. These observations suggest that anthropogenic activities will upset the delicate ecology of these microbes and may lead to massive effect on flora, fauna and human welfare. Therefore there is urgent need to conserve this ecologically delicate and important niche

In conclusion, present study identified the presence of novel groups of bacteria (Fig.6). The current study generated the baseline data of bacterial diversity and composition in sediments of mangroves of west coast of India. The data obtained in the current study clearly indicate that biogeochemical parameters, anthropogenic activities and micro niche play an important role in structuring the bacterial diversity. GIS based prediction suggests that the mangrove microbial communities may decline in future if immediate conservation measures are not implemented.

## Acknowledgement

We thank Prof. Dileep N. Deobagkar for his valuable suggestions.

**Supplementary Information Table S1:** Information of sampling sites

**Supplementary Information Table S2:** Information of chemical parameters of sampling sites

**Supplementary Information Table S3:** Information of bacterial species

**Supplementary Information Table S4:** Alpha Diversity Indices

**Supplementary Information Table S5:** Global beta diversity indices

**Supplementary Information Table S6:** Information of sampling sites used for GIS based prediction

